# Comparison of multi-parallel qPCR and Kato-Katz for detection of soil-transmitted helminth infection among children in rural Bangladesh

**DOI:** 10.1101/629501

**Authors:** Jade Benjamin-Chung, Nils Pilotte, Ayse Ercumen, Jessica R. Grant, Jacqueline R.M.A. Maasch, Andrew M. Gonzalez, Brian P. Abrams, Ashanta C. Ester, Benjamin F. Arnold, Mahbubur Rahman, Rashidul Haque, Alan E. Hubbard, Stephen P. Luby, Steven A. Williams, John M. Colford

## Abstract

An active area of research investigates whether soil-transmitted helminths (STH) can be locally eliminated in endemic settings. In such settings, highly sensitive diagnostics are needed to detect STH infection. We compared double-slide Kato-Katz, the most commonly used copromicroscopic detection method, to multi-parallel quantitative polymerase chain reaction (qPCR) in 2,800 stool samples from children 2-12 years in rural Bangladesh. We estimated the sensitivity and specificity of each diagnostic using Bayesian latent class analysis. Compared to Kato-Katz, STH prevalence using qPCR was almost 3-fold higher for hookworm species and nearly 2-fold higher for *Trichuris trichiura*. *Ascaris lumbricoides* prevalence was lower using qPCR, and 26% of samples classified as *A. lumbricoides* positive by Kato-Katz were negative by qPCR. Amplicon sequencing of the 18S rDNA from 10 samples confirmed that *A. lumbricoides* was absent in samples classified as positive by Kato-Katz and negative by qPCR. The sensitivity of Kato-Katz was 49% for *A. lumbricoides*, 32% for hookworm, and 52% for *T. trichiura*; the sensitivity of qPCR was 79% for *A. lumbricoides*, 93% for hookworm, and 90% for *T. trichiura*. Specificity was ≥ 97% for both tests for all STH except for Kato-Katz for *A. lumbricoides* (specificity = 68%). There were moderate negative, monotonic correlations between qPCR cycle quantification values and eggs per gram quantified by Kato-Katz. While it is widely assumed that Kato-Katz has few false positives, our results indicate otherwise. Our findings suggest that qPCR is more appropriate than Kato-Katz in low intensity infection settings because of its higher sensitivity and specificity.

**Author summary:** Soil-transmitted helminth infections (STH) (e.g., *Ascaris*, hookworm, *Trichuris*) contribute to a large burden of disease among children in low- and middle-income countries. There is increasing interest in implementing large-scale deworming programs to eliminate STH in certain settings. Efforts to monitor whether local elimination has occurred require sensitive diagnostic tests that will not miss positive cases. Kato-Katz, a microscopy-based diagnostic test, has commonly been used to identify STH eggs in stool, but in settings where infection intensity is low, this method frequently misses positive samples because it requires visual identification of small numbers of eggs, and eggs may degrade prior to visualization. Quantitative polymerase chain reaction (qPCR) is a molecular diagnostic method that may miss fewer infections because it identifies STH DNA in stool, which can be detected in very small quantities and is less likely to degrade. This study compared the performance of Kato-Katz and qPCR using 2,800 stool samples from children aged 2-12 years in rural Bangladesh. qPCR detected substantially more hookworm and *Trichuris* infections than Kato-Katz. 26% of samples were classified as *Ascaris* positive by Kato-Katz and negative by qPCR. We conclude that qPCR is a more appropriate diagnostic method than Kato-Katz in low infection intensity settings.

## Introduction

Soil-transmitted helminths (STH) infect an estimated 1.45 billion individuals, almost one fifth of the global population (1). These infections contribute to a substantial burden of disability and disease, particularly for children (2). To control the burden of STH infection, mass drug administration programs deliver presumptive chemotherapy to populations at risk, such as school children. Such programs have expanded dramatically following the London Declaration in 2012, and in 2010 approximately 314 million children – 31% of children at risk – received deworming medication for soil-transmitted helminths (3). Mass drug administration programs have yielded large reductions in STH infection prevalence and intensity to date (4). While the World Health Organization has set goals for STH focused on morbidity control (3), there is an active body of research focused on whether local elimination of STH is possible in endemic settings through community-wide mass deworming administration (5–8). As STH infection intensity and prevalence wane, increasingly sensitive diagnostic methods are required to determine whether STH transmission has been interrupted and when interventions can be scaled back or discontinued (9).

Diagnostic methods for STH include copromicroscopic methods, such as the Kato-Katz method (10,11), formol-ether concentration (12), and FLOTAC (13). Of these, Kato-Katz is the most commonly used diagnostic method for STH surveillance because it is inexpensive and relatively easy to perform in low-resource settings (14). A significant limitation of this method is that samples must be evaluated within an hour of preparation in order to detect fragile hookworm ova. In addition, *Necator americanus* and *Ancylostoma duodenale* cannot be distinguished using Kato-Katz because of their morphological similarities. Because egg excretion is highly variable over time (15), Kato-Katz has improved sensitivity when performed on multiple samples from defecation events on different days, but such samples are often logistically prohibitive to collect. Double-slide Kato-Katz examines stool using two separate aliquots from a single defecation event prepared as separate slides in order to help reduce the variability and increase sensitivity. Furthermore, a meta-analysis estimated the sensitivity of double-slide Kato-Katz to be 64.6% for *Ascaris lumbricoides*, 84.8% for *Trichuris trichiura*, and 63.0% for hookworm; however, in low-intensity settings sensitivity was lower: 55.2% for *A. lumbricoides*, 79.8% for *T. trichiura*, and 52.6% for hookworm (16). Alternative copromicroscopic STH diagnostic methods, such as formol-ether concentration, FLOTAC, mini-FLOTAC, and the McMaster method also have moderate to low sensitivity for different types of STH in low-intensity settings (16).

Multiplex (17,18) and multi-parallel (19,20) quantitative polymerase chain reaction (qPCR) assays have been developed to detect STH DNA in stool. These molecular methods are likely to be more sensitive than microscopy because DNA can still be detected in stool samples when ova have disintegrated and can no longer be visualized using copromicroscopy. Multiplex and multi-parallel assays allow for detection of multiple helminths from a single stool sample, and growing evidence suggests that these methods may be used to estimate infection intensity (18,21,22). Five studies in Ghana, Kenya, Argentina, and Ecuador have compared diagnostic performance of copromicroscopic methods and qPCR and reported that compared to Kato-Katz, qPCR had higher detection rates (19), sensitivity, and specificity (17,23,21,22).

The greater sensitivity, specificity, and precision of qPCR relative to Kato-Katz makes it an attractive diagnostic for monitoring the successes of STH control programs deploying mass deworming since infection intensity tends to decline after several years of intervention (24). As a population reaches an STH transmission breakpoint, worm burdens decrease, and the comparatively low sensitivity of Kato-Katz may preclude its use in monitoring STH prevalence. In addition, a growing body of work suggests that qPCR may also be used to estimate infection intensity; one study found that qPCR is approximately 3.6 times more precise than Kato-Katz in estimating *A. lumbricoides* egg intensity (25). Such additional precision could increase the statistical power available to detect whether the STH prevalence meets the threshold for elimination.

The objectives of this study were to estimate the prevalence of STH infection and infection intensity using Kato-Katz and multi-parallel qPCR and to estimate the sensitivity and specificity of each method using archived samples collected from children aged 2-12 years in rural Bangladesh.

## Methods

### Study population

This study analyzed archived stool samples that were collected from participants in the WASH Benefits Bangladesh trial between May 2015 and May 2016 (Clinicaltrials.gov NCT01590095) (26,27). The trial was implemented in Gazipur, Mymensingh, Tangail and Kishoreganj districts of Bangladesh. Biannual school-based mass drug administration with mebendazole had been implemented nationally in Bangladesh since 2008 and was ongoing in the study area during the trial. The trial delivered interventions to improve water, sanitation, handwashing, and nutrition and included single and combined intervention arms. Full details of interventions and user adherence have been published elsewhere (27,28). This study randomly selected 2,800 stool specimens collected from children aged 22 months to 12 years of age (mean = 57 months) enrolled in the individual improved water, sanitation, handwashing, and combined water+sanitation+handwashing (WSH) arms. This sample size was chosen in order to estimate sensitivity with precision of +/− 5% and 80% statistical power; we assumed STH prevalence estimates from a prior study in rural Bangladesh (29) and Kato-Katz sensitivity and specificity estimates from a prior meta-analysis of STH diagnostic accuracy (16).

### Stool collection and storage

Field staff provided study participants’ primary caregivers with sterile containers and requested that they collect stool from the child’s defecation event the next morning. Field staff returned to the house at least three times to attempt to collect stool containers. All study participants received a single dose of albendazole following stool collection. Field staff transported stool specimens on ice to the field laboratory, where 1 g of stool was archived in 1 ml of 100% ethanol. Specimens were stored at −20° C prior to relocation to a −80° C freezer. In the WASH Benefits parasites sub-study, the median time that elapsed between the reported time of defecation and when the stool was first placed on ice was 2.4 hours (range: 0.03 to 18 hours), and the median time that elapsed between the date of specimen collection and the date when specimens were stored in a −80° C freezer was 11 days (range: 1 to 330 days).

### Kato-Katz procedures

Technicians were trained in Kato-Katz using the Vestergaard Frandsen protocol at the icddr,b parasitology laboratory. On the day of stool collection, trained technicians performed double-slide Kato-Katz (10,11) on fresh stool within 30 minutes of preparing each sample. For quality assurance, 10% of slides were evaluated independently by two technicians, and 5% were evaluated by an experienced parasitologist. The quality assurance results have previously been published elsewhere (28). In brief, agreement between laboratory technicians was high (Kappa statistic > 0.99 for each STH). The Kappa statistic for agreement between laboratory technicians and experienced parasitologists was 0.92 for *A. lumbricoides*, 0.20 for hookworm, and 0.86 for *T. trichiura*. The agreement is likely lower for hookworm because experienced parasitologists examined slides up to a few days after laboratory technicians, and hookworm ova may have begun to disintegrate by that time. Samples in which at least one STH egg was visualized during Kato-Katz were classified as positive. Kato-Katz technicians did not have access to any clinical information about study participants.

### DNA isolation and qPCR procedures

Preserved stool specimens were shipped to Smith College in Northampton, MA, United States for qPCR analyses. Prior to shipment, ethanol was evaporated from all samples for compliance with shipping regulations, and samples were shipped on dry ice. Upon receipt at Smith College, all samples were stored at −20 °C until DNA extraction. DNA was extracted using the FastDNA Spin Kit for Soil (MP Biomedicals, Solon, OH) utilizing a modified version of the previously published methodology (19). An internal amplification control (IAC) plasmid was employed during the extraction of each sample to ensure successful isolation, and adequate recovery of DNA (30). Utilizing previously described reaction methods (31), any sample which failed to produce a positive qPCR result for one or more of the IAC replicates underwent re-extraction. Similarly, following the completion of all extractions, the mean quantitation cycle (Cq) value for the IAC results from all samples was calculated, as was the standard deviation of the mean. Any sample with an individual IAC result determined to be 3 or more standard deviations greater than the mean underwent re-extraction, as such a recovery was deemed suboptimal. Following re-extraction, only the extracted DNA sample producing a positive IAC result within the defined Cq range (not the previously conducted invalid extract) was used for experimental testing.

Experimental qPCR reactions were performed using previously published multi-parallel assays targeting non-coding repetitive sequences to detect *Ascaris lumbricoides* (32)*, Trichuris trichiura, Strongyloides stercoralis, Necator americanus, Ancylostoma duodenale* (20) *and Ancylostoma ceylanicum* (33). Laboratory staff were blinded to Kato-Katz results for each sample during the initial qPCR analyses. However, upon completion of initial testing, all samples determined to be Kato-Katz positive but qPCR negative for *A. lumbricoides* were tested using a second qPCR assay targeting an unrelated, ribosomal DNA sequence in order to validate the negative qPCR result and confirm the absence of *A. lumbricoides* DNA (20,34).

For each assay, all samples were tested in replicate reactions and samples were classified as positive when amplification occurred in both reactions with a Cq value of <40, consistent with prior studies (35). Samples which produced Cq values ≥40 in both replicate reactions or failed to amplify in both replicate reactions were classified as negative. Samples that were positive in one of two replicate reactions were re-tested in duplicate and classified as positive if the Cq value was <40 in at least one of two re-test reactions. In all cases of re-testing, the Cq value reported as the sample’s final result was the re-test Cq value.

In addition to the IAC control, all experimental qPCR reaction plates were accompanied by the testing of a titration of the appropriate assay’s control plasmid. Plasmid controls for each target sequence were prepared as previously described (32). Each plasmid contained a single copy of the corresponding assay’s target sequence and 20 pg, 200 fg and 2 fg of plasmid were tested in duplicate reactions. Following the completion of all testing, the mean Cq value, across all experimental plates, for each plasmid concentration was determined. Any plate which produced a Cq value for any concentration of plasmid that was 3 or more standard deviations greater than the mean calculated for all plates was retested in its entirety, and all results from that plate were considered to be invalid. Only results from valid plates were reported.

### Sequencing

A subset of 10 samples was selected to undergo amplicon sequencing-based analysis. To facilitate the selection of samples, all samples that were Kato-Katz positive for the presence of *A. lumbricoides* but qPCR negative for *A. lumbricoides* were identified. From this list, a convenience sample (n=7) was then selected, such that two of these samples would contain eggs per gram (epg) counts characteristic of moderate-intensity infections as determined by World Health Organization guidelines, while five samples would contain egg counts that were characteristic of light intensity infections (36). All seven of the Kato-Katz positive samples were negative by qPCR using both *A. lumbricoides* assays. Additionally, three Kato-Katz negative samples were chosen for inclusion in this panel. Two of these samples were selected due to their status as *A. lumbricoides-*positive as determined by qPCR analysis, while a single sample was selected that was both Kato-Katz negative and qPCR negative to serve as an uninfected control.

Samples were prepared for sequencing using a modified version of the Earth Microbiome Project’s 18S Illumina Amplicon Protocol available at: http://www.earthmicrobiome.org/protocols-and-standards/18s/. This protocol utilizes primers targeting the variable region 9 (V9) of the eukaryotic small subunit (SSU) rDNA (37). Briefly, the targets within each sample were amplified in triplicate 25 µl reactions using a uniquely barcoded reverse primer coupled with a conserved forward primer (Table 1). A “mammalian blocking” primer, based on a previously described strategy, was also included in all reactions to minimize the prevalence of mammal-derived amplicons (38). All reactions contained 10 µl of Platinum Hot Start PCR Master Mix (2X) (ThermoFisher, Waltham, MA), 0.2 µM forward and reverse primers, and 1.6 µM “mammalian blocking” primer (Integrated DNA Technologies, Coralville, IA). Cycling consisted of an initial denaturation step of 94 °C for 3 minutes, followed by 35 cycles of 94 °C for 45 seconds, 65 °C for 15 seconds, 57 °C for 30 seconds, and 72 °C for 90 seconds. Samples then underwent a final extension step at 72 °C for 10 minutes. Following amplification, triplicate reactions were combined, and pooled products were run on a 1.5% agarose gel to confirm the presence of a band of the expected size (~260 bp). 240 ng of each pooled product was then combined in preparation for sequencing, and this library was purified using the ZR-96 Clean & Concentrator purification kit (Zymo Research, Irvine, CA). The purified library was diluted to a 9 pM concentration, and 30% PhiX was added to improve diversity. Sequencing then occurred on the Illumina MiSeq platform, using a MiSeq Reagent Kit v2 (300-cycles) (Illumina, San Diego, CA).

**Table 1.**
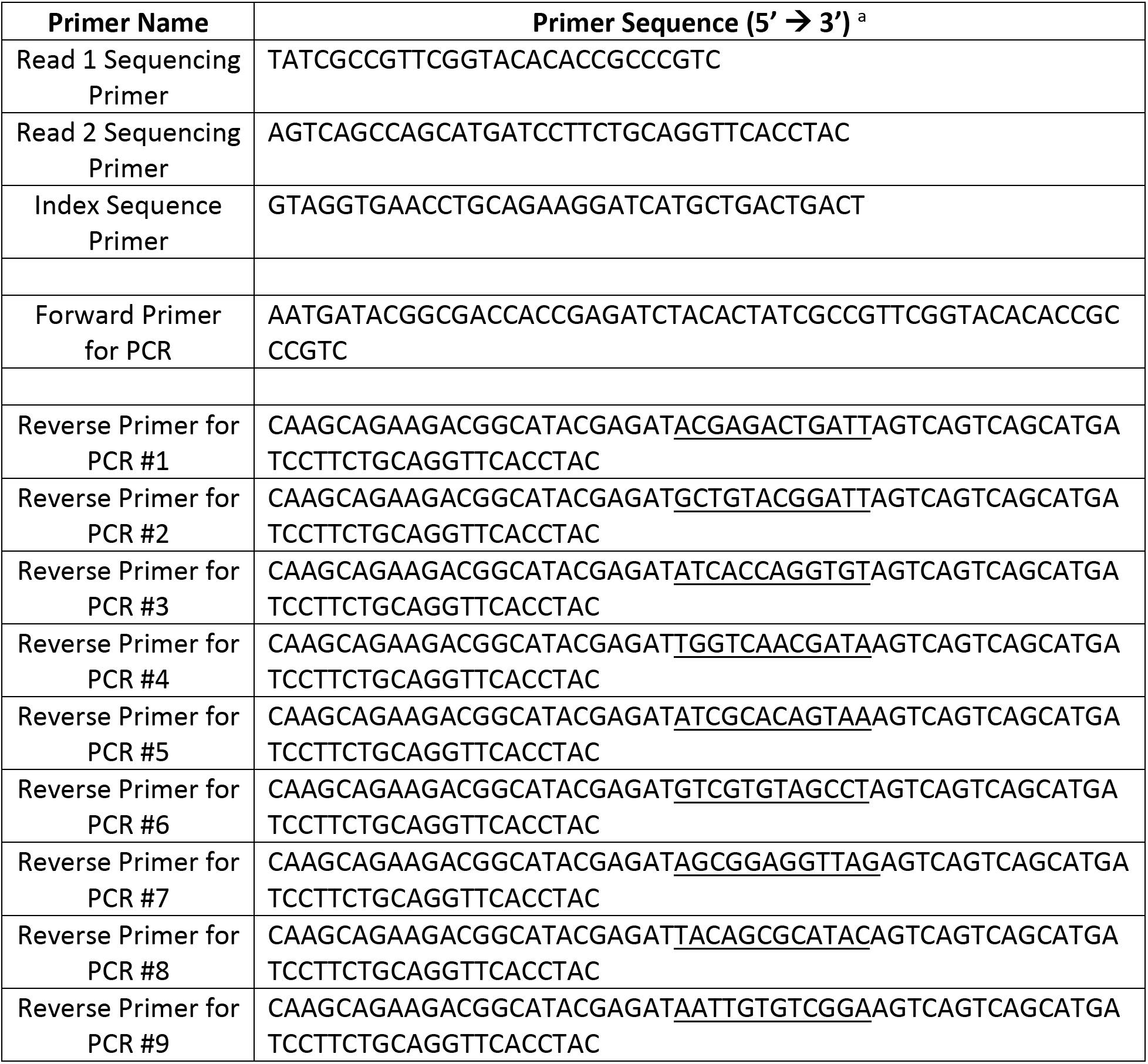
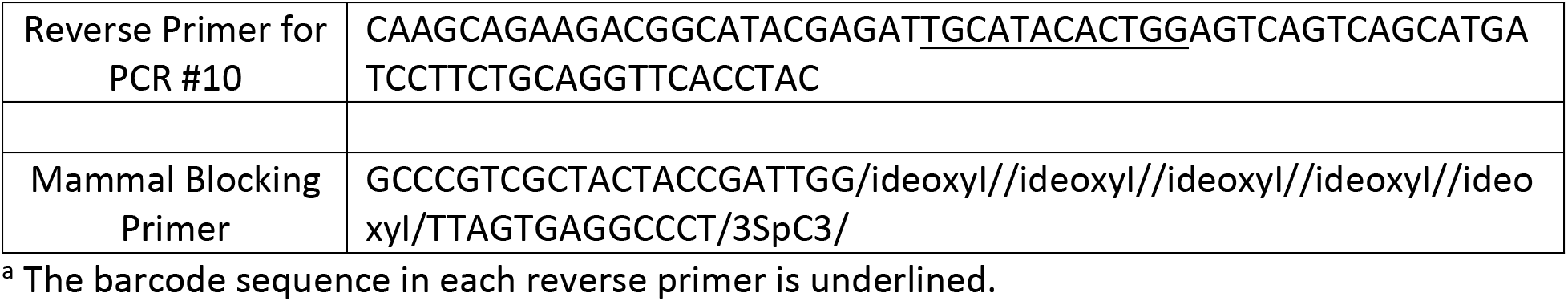
Primer sequences used for 18S SSU sequencing.

Sequencing results were analyzed using the Quantitative Insights into Microbial Ecology 2 (QIIME2) pipeline (39). Briefly, all interlaced read pairs which did not contain a valid barcode were filtered out and all read pairs whose best match was to bacterial 16S ribosomal sequence were also removed. This resulted in a list of reads exclusively matching eukaryotic 18S sequence. Reads with 99% or greater identity were then classified into operational taxonomic units (OTUs) and consensus sequences were built. Taxonomy was then assigned to all OTUs that had ≥ 95% identity to sequences within the Silva release 132 QIIME-compatible database.

### Statistical methods

All statistical analyses pooled across intervention arms to ensure sufficient statistical power. Unless otherwise specified, analyses were conducted in R version 3.4.3. Data and replication scripts are available through the Open Science Framework (https://osf.io/agk6w/). The STARD Checklist for this study is included in S1 Appendix.

#### Prevalence and correlation between eggs per gram and Cq values

We estimated the prevalence of STH infection under each diagnostic method with robust sandwich standard errors to account for clustering at the village level (40). We calculated the agreement between prevalence estimates from Kato-Katz and qPCR using a kappa statistic. We tested whether prevalence differed between the two diagnostic methods using the Wilcoxon matched-pairs signed rank test that conditioned on randomized block in the original trial (26,41). To classify infection intensity, we used the World Health Organization cutoffs (moderate/high intensity infections defined as ≥5,000 eggs/gram for Ascaris, ≥1,000 eggs/gram for Trichuris, and ≥2,000 eggs/gram for hookworm) (36). To assess the correlation between epg estimated by Kato-Katz and Cq value estimated by qPCR, we estimated Kendall’s tau and calculated two-sided p-values using a bootstrap with 1,000 replicates that resampled blocks to account for the study’s cluster-randomized design using Stata version 14.2. We compared mean Cq values within levels of infection intensity using a paired t-test paired within randomized blocks.

#### Estimation of sensitivity and specificity

A challenge in studies assessing accuracy of STH diagnostics is the lack of a gold standard diagnostic method (42). We thus used two different approaches to estimate sensitivity and specificity of each method, consistent with prior studies. We created a reference using both detection methods and defined positivity as qPCR Cq values < 40 or at least one egg detected by Kato-Katz. We also used Bayesian latent class analysis, which defines the true prevalence, sensitivity, and specificity as latent variables that are estimated simultaneously from the data and assumes no gold standard (16,22,43–45). We defined a model that accounted for covariance between diagnostic methods since both methods are dependent upon the same underlying biological process (43) (S2 Appendix). As has been done in prior studies, we used a highly informative prior for specificity to allow for identifiability (16,22,46,47) since prior studies have reported high specificity values for Kato-Katz (48,49) and qPCR (19,50) (S3 Appendix Table 1). For *A. lumbricoides*, we chose to specify a non-informative prior for Kato-Katz sensitivity and specificity and a less informative prior for qPCR sensitivity because of discordant test results (summarized below). To assess the impact of these priors on estimates, we conducted a sensitivity analysis using alternative, less conservative priors (S3 Appendix Table 2). To estimate parameters, we used Monte Carlo simulation with 4 chains and 100,000 iterations, recording every 10^th^ result. We computed the mean and 2.5 and 97.5 percentiles of the posterior distribution using Gibbs sampling in WinBUGS version 14 (51).

**Table 2.** Soil-transmitted helminth prevalence and intensity.

|  | Kato-Katz (N=2800) |  | qPCR (N=2799) |  |
| --- | --- | --- | --- | --- |
|  | Prevalence (95% CI) | Geometric mean <sup>a</sup> EPG (95% CI) | Prevalence (95% CI) | Median Cq value in positive stool samples (range) |
| *Ascaris lumbricoides* | 37.0 (34.7, 39.3) | 5.14 (4.42, 5.96) | 23.3 (20.9, 25.7) | 17.4 (4.4, 39.6) |
| Hookworm | 7.5 (6.3, 8.7) | 0.43 (0.35, 0.53) | 21.4 (19.4, 23.3) | -- |
| *Necator americanus* | -- | -- | 18.4 (16.5, 20.3) | 20.8 (13.9, 35.5) |
| *Ancylostoma ceylanicum* | -- | -- | 3.8 (2.9, 4.6) | 23.6 (14.4, 37.8) |
| *Ancylostoma duodenale* | -- | -- | 0.0 (-0.01, 0.10) | 27.2 (27.2, 27.2) |
| *Trichuris trichiura* | 7.0 (5.6, 8.3) | 0.40 (0.30, 0.50) | 12.3 (10.3, 14.2) | 27.7 (21.4, 40.0) |
| *Strongyloides stercoralis* | -- | -- | 0.6 (0.3, 0.9) | 24.6 (19.1, 31.8) |

<sup>a</sup> Includes both negative and positive samples.

### Ethical statement

The study protocol was approved by the Ethical Review Committee at The International Centre for Diarrhoeal Disease Research, Bangladesh (PR-14105), the Committee for the Protection of Human Subjects at the University of California, Berkeley (2014-08-6658), and the institutional review board at Stanford University (27864). Prior to enrollment, all adult subjects provided written informed consent. Parents or guardians of children provided written informed consent on behalf of children.

## Results

We analyzed stool samples from 2,800 children using both Kato-Katz and qPCR. 51% of children were female, and 66% of children’s caregivers reported that they had received deworming in the past 6 months (S4 Appendix). We excluded one sample’s qPCR results from statistical analyses for all species because the internal amplification control was >3 standard deviations from the mean upon initial testing and following re-extraction.

The detection rate was higher for qPCR than Kato-Katz for hookworm and *T. trichiura* but lower for *A. lumbricoides* (Table 2, Fig 1). The prevalence of *A. lumbricoides* was 37.0% (95% CI 34.7%, 39.3%) using Kato-Katz and 23.3% (95% CI 20.9%, 25.7%) using qPCR. The prevalence of hookworm was 7.5% (95% CI 6.3%, 8.7%) and 21.4% (95% CI 19.4%, 23.3%) for any hookworm species using qPCR. Using qPCR the prevalence of *Necator americanus* was 18.4% (95% CI 16.5%, 20.3%), and the prevalence of *Ancylostoma ceylanicum* was 3.8% (95% CI 2.9%, 4.6%); only one out of 2,799 samples tested positive for *Ancylostoma duodenale*. For *T. trichiura*, the prevalence was 7.0% (95% 5.6%, 8.3%) by Kato-Katz and 12.3% (95% CI 10.3%, 14.2%) by qPCR. Under Kato-Katz, 7% of children were infected with more than one STH, and under qPCR it was 13% (Fig 2). The prevalence of moderate-to-heavy intensity infection under Kato-Katz was 4% for *A. lumbricoides* and less than 1% for hookworm and *T. trichiura*. Using Kato-Katz, the geometric mean of epg was 5.14 for *A. lumbricoides* (134 among positives), 0.43 for hookworm (125 among positives), and 0.40 for *T. trichiura* (124 among positives).

**Fig 1.**
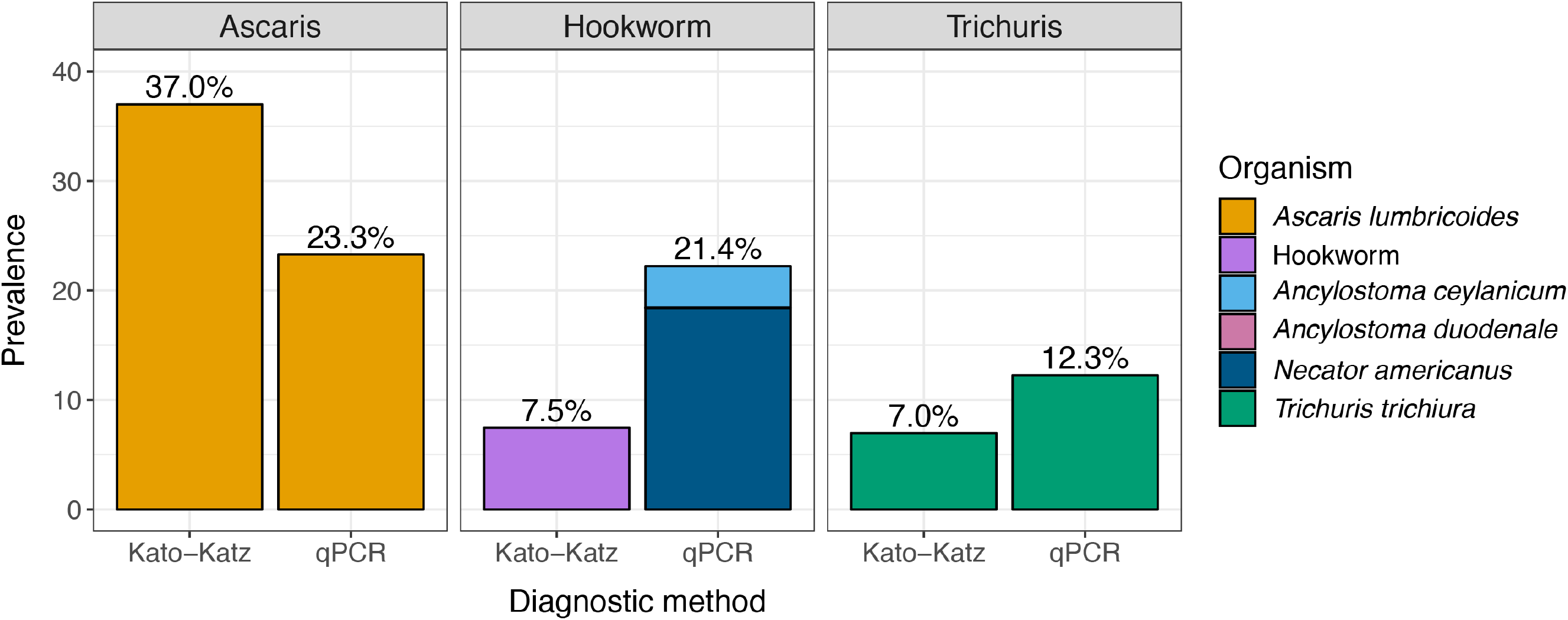
Soil-transmitted helminth prevalence by Kato-Katz and qPCR. Prevalence was estimated from stool samples collected from children aged 2-12 years in rural Bangladesh (Kato-Katz: N=2,800; qPCR N=2,799).

**Fig 2.**
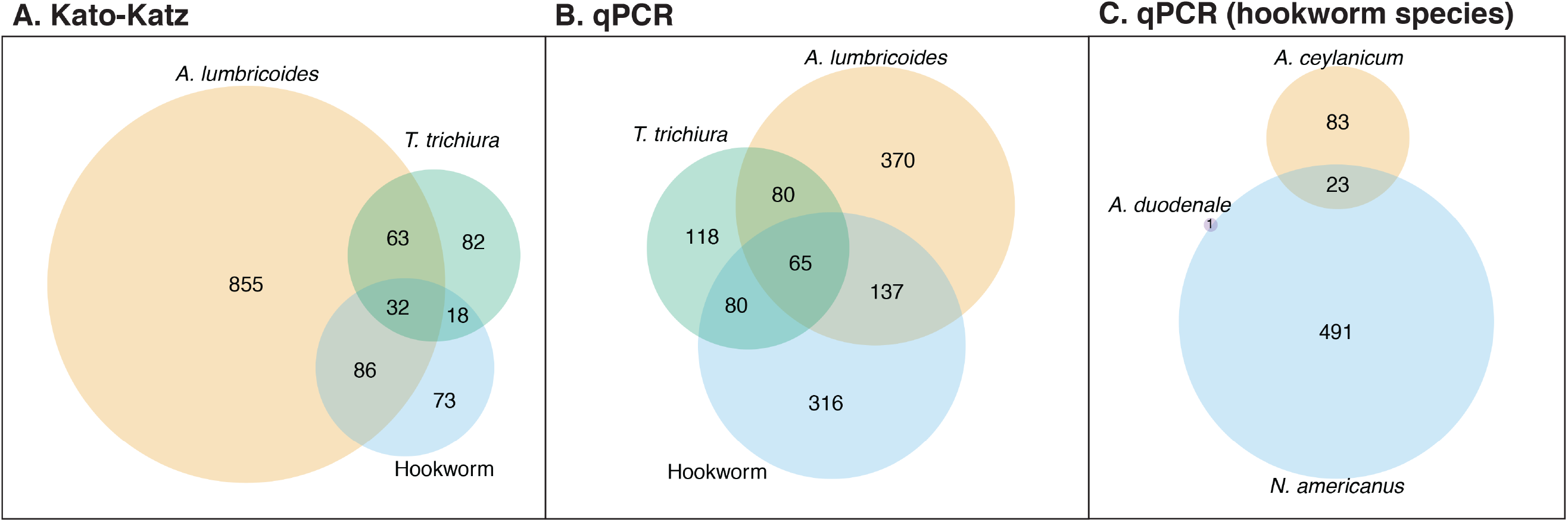
Venn diagram of co-infections detected by Kato-Katz and qPCR. Soil-transmitted helminth ova or DNA were detected in stool samples collected from children aged 2-12 years in rural Bangladesh using Kato-Katz (N=2,800) or multi-parallel qPCR (N=2,799).

Concordance between Kato-Katz and qPCR was higher for hookworm and *Trichuris* than for *Ascaris*. The p-value for permutation tests assessing whether the prevalence estimated by the two detection methods differed was <0.001 for each STH (Table 3). 6-14% of samples classified as negative for the different species by Kato-Katz were positive using qPCR. 26% of samples classified as positive for *A. lumbricoides* by Kato-Katz were negative using qPCR. All samples determined to be Kato-Katz positive for *A. lumbricoides* but negative by qPCR analysis targeting the non-coding repeat also failed to produce a positive result when tested with the ITS-targeting assay.

**Table 3.** Classification of qPCR and Kato-Katz for each type of STH.

|  | qPCR + | qPCR - | Kappa statistic (p-value) |
| --- | --- | --- | --- |
| *A. lumbricoides* |  |  |  |
| Kato-Katz + | 319 (11%) | 716 (26%) | 0.13 (<0.001) |
| Kato-Katz - | 333 (12%) | 1431 (51%) |  |
| Hookworm |  |  |  |
| Kato-Katz + | 195 (7%) | 14 (1%) | 0.42 (<0.001) |
| Kato-Katz - | 403 (14%) | 2187 (78%) |  |
| *T. trichiura* |  |  |  |
| Kato-Katz + | 167 (6%) | 28 (1%) | 0.58 (<0.001) |
| Kato-Katz - | 176 (6%) | 2428 (87%) |  |

Using Bayesian latent class models, the estimated true prevalence was 27% (95% Bayesian Credible Interval (BCI) 20%, 37%) for *A. lumbricoides*, 20% (95% BCI 17%, 24%) for hookworm, and 11% (95% BCI 8%, 14%) for *T. trichiura*. For Kato-Katz, the sensitivity was 49% (95% BCI 34%, 64%) for *A. lumbricoides*, 32% (95% BCI 22%, 41%) for hookworm, and 52% (95% BCI 33%, 71%) for *T. trichiura*, and the specificity was 68% (95% BCI 61%, 77%) for *A. lumbricoides*, 99% (95% BCI 96%, 100%) for hookworm, and 98% (95% BCI 96%, 100%) for *T. trichiura* (Table 4). For qPCR, the sensitivity was 79% (95% BCI 61%, 99%) for *A. lumbricoides*, 93% (95% BCI 82%, 100%) for hookworm, and 90% (95% BCI 81%, 100%) for *T. trichiura*, and the specificity was 97% (95% BCI 95%, 100%) for all three STH. The sensitivity analysis for *A. lumbricoides* using alternative less conservative priors produced similar estimates for Kato-Katz and higher sensitivity for qPCR; when more informative priors were used for both Kato-Katz and qPCR, the estimated sensitivity of qPCR was 90 (95% BCI 80, 99) (S3 Appendix Table 3). Pooling both methods as the gold standard, the sensitivity and specificity of Kato-Katz was similar to those estimated from the Bayesian model, and for qPCR the sensitivity of hookworm and *T. trichiura* was higher but the specificity was lower than estimated by the Bayesian model.

**Table 4.** Estimated prevalence, sensitivity, and specificity of each diagnostic method using Bayesian latent class models.

|  | Kato-Katz |  | qPCR |  |
| --- | --- | --- | --- | --- |
|  | Sensitivity (%)<br>(95% BCI) | Specificity (%)<br>(95% BCI) | Sensitivity (%)<br>(95% BCI) | Specificity (%)<br>(95% BCI) |
| **Bayesian latent class model** |  |  |  |  |
| *Ascaris lumbricoides* | 49 (34, 64) | 68 (61, 77) | 79 (61, 99) | 97 (95, 100) |
| Hookworm<sup>a</sup> | 32 (22, 41) | 99 (96, 100) | 93 (82, 100) | 97 (95, 100) |
| *Trichuris trichiura* | 52 (33, 71) | 98 (96, 100) | 90 (81, 100) | 97 (95, 100) |
| **Any positive test (sensitivity) or any negative test (specificity) under Kato-Katz or qPCR** |  |  |  |  |
|  | Sensitivity (%)<br>(95% CI) | Specificity (%)<br>(95% CI) | Sensitivity (%)<br>(95% CI) | Specificity (%)<br>(95% CI) |
| *Ascaris lumbricoides*<sup>b</sup> | 49 (45, 53) | 67 (64, 69) | -- | -- |
| Hookworm<sup>a</sup> | 34 (30, 38) | 99 (99, 100) | 98 (97, 99) | 85 (83, 86) |
| *Trichuris trichiura* | 53 (47, 58) | 99 (99, 99) | 92 (90, 95) | 93 (92, 94) |

95% BCI: 95% Bayesian credible interval
95% CI: 95% confidence interval
<sup>a</sup> *Necator americanus*, *Ancylostoma duodenale*, and *Ancylostoma ceylanicum* combined
<sup>b</sup> Due to the high discordance between qPCR and Kato-Katz for *A. lumbricoides*, we estimated sensitivity and specificity using qPCR alone as the gold standard.

For each STH, there was a negative, monotonic relationship between qPCR Cq values and epg estimated using Kato-Katz (Fig 3). The Kendall’s tau comparing these two quantities was - 0.442 for *A. lumbricoides*, −0.346 for *N. americanus*, −0.266 for *A. ceylanicum*, and −0.248 for *T. trichiura* (for each, the p-value was <0.001); we compared *N. americanus* and *A. ceylanicum* to any hookworm species detected by Kato-Katz. We also examined the distribution of Cq values by Kato-Katz infection intensity status for *A. lumbricoides* (very few children had moderate-to-heavy intensity hookworm or *T. trichiura* infections using Kato-Katz) (Fig 4). The distribution of Cq values was higher for children with light intensity infections compared to those with moderate-to-heavy infections. The median Cq value was 20 (range: 16-29) among children with moderate to heavy *A. lumbricoides* infection and 25 (range: 20-36) among those with light intensity infection (t-test p-value < 0.001).

**Fig 3.**
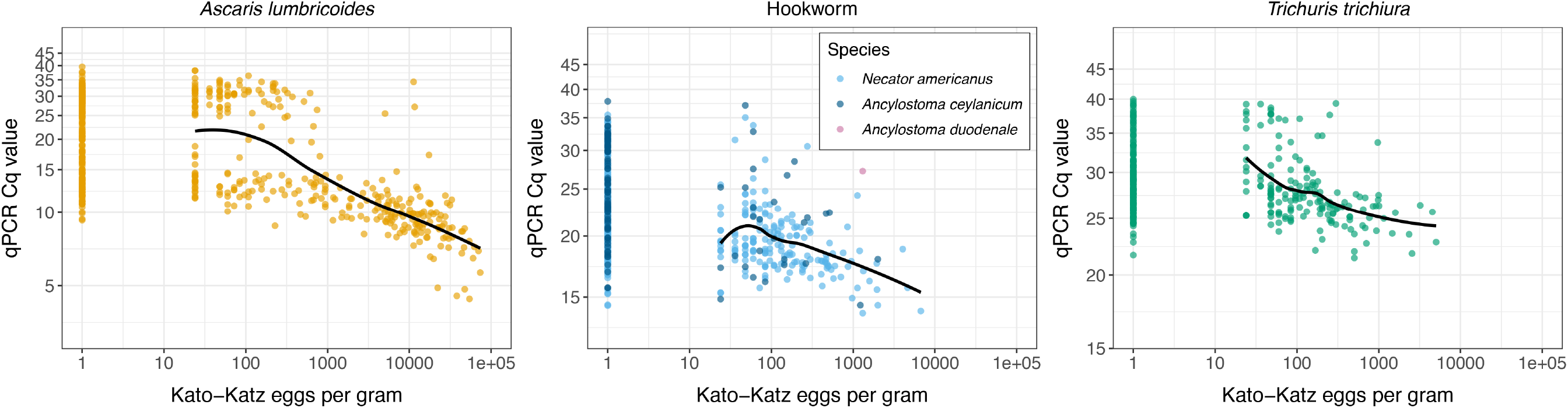
Relationship between Cq values measured by qPCR and eggs per gram estimated using Kato-Katz. Soil-transmitted helminth ova or DNA were detected in stool samples collected from children aged 2-12 years in rural Bangladesh using Kato-Katz (N=2,800) or multi-parallel qPCR (N=2,799). The Kendall’s tau comparing eggs per gram and Cq values was −0.442 for *A. lumbricoides*, −0.346 for *N. americanus*, −0.266 for *A. ceylanicum*, and −0.248 for *T. trichiura* (for each, the p-value was <0.001).

**Fig 4.**
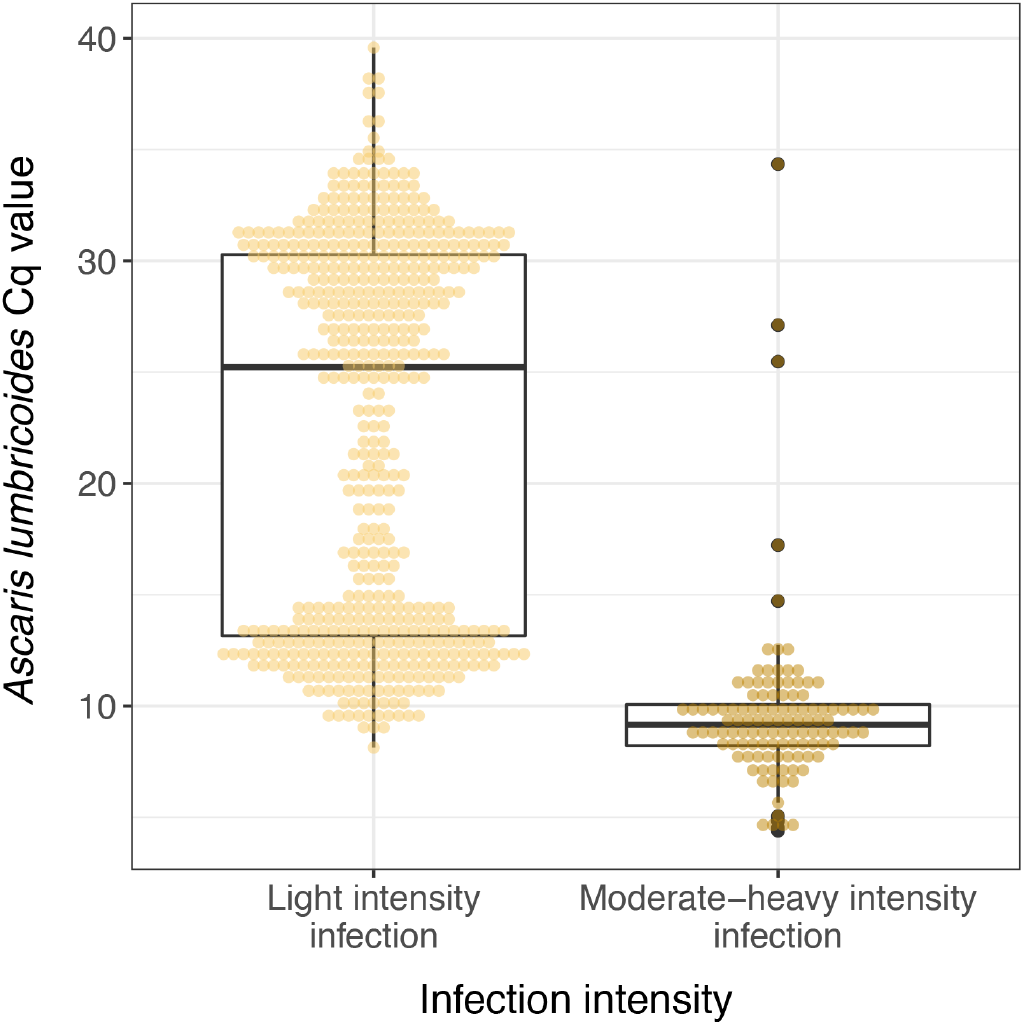
Distribution of *A. lumbricoides* Cq values within infection intensity categories using Kato-Katz. Light intensity infection was defined as < 5,000 eggs per gram from Kato-Katz, and moderate/heavy intensity infections were defined as ≥5,000 eggs per gram per the World Health Organization definition.

In the sequencing analyses of 10 samples, we detected *Enterobius vermicularis* in one sample that was Kato-Katz negative and qPCR positive, one sample that was Kato-Katz positive and qPCR negative, and one sample that was negative under both methods (S5 Appendix). In addition, we identified *Giardia intestinalis* in 3 samples that were Kato-Katz positive and qPCR negative and one sample that was negative using both methods; we detected *Dientamoeba fragilis* in 3 samples that were Kato-Katz positive and qPCR negative, one sample that was Kato-Katz negative and qPCR positive, and one sample that was negative using both methods. Analysis using the QIIME2 pipeline failed to identify any reads mapping to *Ascaris* in all seven Kato-Katz positive, qPCR negative samples. *A. lumbricoides* was detected in one of the two samples that were Kato-Katz negative and qPCR positive. However, this analysis failed to identify *A. lumbricoides* in the second sample. Because of this discrepancy, the *A. lumbricoides-* derived consensus sequence of the reads from the *A. lumbricoides-*positive sample was compared, using BLAST, to all other reads from the entire sample set. This resulted in the identification of a small number of *A. lumbricoides-*derived reads (n = 5, identity ≥ 97%) within the second Kato-Katz negative, qPCR positive sample, while no *A. lumbricoides* reads were found within the pool of reads mapping to any of the other samples. As only a single read within this Kato-Katz negative, qPCR positive sample had 100% identity with the *A. lumbricoides-*derived consensus sequence used as the query sequence in the BLAST analysis, QIIME2’s failure to generate an OTU from these reads follows logically, as it is likely that the 99% similarity threshold was not met for any two reads mapping to *Ascaris* from this sample. However, the presence of *A. lumbricoides-*derived reads provides further support for the existence of *A. lumbricoides* DNA within the sample and speaks to the sensitivity of the qPCR assays used during the analysis.

## Discussion

This study compared the performance of Kato-Katz and qPCR for detecting STH infections in a large sample of 2,800 children in a setting in rural Bangladesh with biannual mass deworming administration. Consistent with prior studies in low STH infection intensity settings, using qPCR led to substantial increases in hookworm detection and moderate increases in *T. trichiura* detection compared to Kato-Katz (17,19,21,22,42). Poorer performance of Kato-Katz, particularly for hookworm, is likely due to rapid disintegration of hookworm ova following stool collection (42,52). However, we found that the detection rate for *A. lumbricoides* was 14% lower for qPCR than Kato-Katz. These unexpected findings call into question the common assumption that Kato-Katz yields few false positives (16,22,46,47). The quality control assessment found minimal misclassification of *A. lumbricoides* between laboratory technicians and expert parasitologists (53), suggesting that Kato-Katz was performed with typical quality. Yet, experimental testing with two qPCR assays utilizing independent targets as well as sequencing analysis of a subset of samples confirmed that samples classified as *A. lumbricoides* negative by qPCR indeed contained no genomic targets for *A. lumbricoides*.

Misclassification of STH ova has been previously reported: *Capillaria philippinensis* or *Capillaria hepatica* ova can be mistaken for *A. lumbricoides* or *T. trichiura* ova (54,55) and *Trichostrongylus spp*. or *Schistosoma* ova can resemble hookworm ova (56,57). Though we did identify other enteropathogens (*Giardia intestinalis* and *Dientamoeba fragilis)* when sequencing a subset of samples, in stool, these parasites do not share morphological features with most stages of *A. lumbricoides* (58). We believe that the substance mistaken for *A. lumbricoides* in Kato-Katz was most likely plant material, pollen grains, or fungal spores, all of which can resemble certain stages of *A. lumbricoides* (59). Some of the sequenced stool samples contained DNA of flowering plants of the division *Magnoliophyta*, which produce pollen, and fungi of the genera *Saccharomyces* and *Candida*, which are commonly found in healthy intestinal microbiota (60) (S5 Appendix). We cannot definitively identify what organism or substance was mistaken for *A. lumbricoides* because the original slides used in Kato-Katz were not preserved. Nevertheless, our findings underscore the advantage of using molecular methods over copromicroscopy since molecular methods do not rely on visualization of stool contents.

Overall, we found that sensitivity was lower for both double-slide Kato-Katz and qPCR than in prior studies (16,22). A challenge in quantifying the accuracy of STH diagnostics is the lack of a gold standard measure of infection (9). Though we used Bayesian latent class analysis, which assumes no gold standard, our estimates of sensitivity and specificity from this method should be interpreted with caution. In studies evaluating fewer than four diagnostic tests, parameters are not identifiable, and as a result parameter estimates are highly dependent upon the specified prior distribution and the disease prevalence (43). Thus, our estimates of true prevalence, sensitivity and specificity using this method are sensitive to our modeling assumptions. For hookworm and *T. trichiura* there was internal validity between results from the Bayesian latent class analysis and an analysis pooling both methods as the reference. However, because of the unexpectedly high degree of discordance between Kato-Katz and qPCR for *A. lumbricoides*, we used less informative priors for the specificity of Kato-Katz and the sensitivity of qPCR. Our estimate of sensitivity of qPCR for detecting *A. lumbricoides* using Bayesian latent class analysis was lower than a previous estimate for multi-parallel qPCR using the same method (22). Using less informative priors may have reduced the identifiability of our model and contributed to differing sensitivity and specificity estimates in this study. However, our sensitivity analysis using more informative priors for *A. lumbricoides* produced estimates that were similar to our primary analysis and/or closer to previously published estimates (S3 Appendix Tables 2–3).

In the absence of intact eggs, Kato-Katz will produce a negative result, even if STH DNA is present in stool. An advantage of qPCR is that it can detect DNA quantities less than the amount derived from a single egg – an important feature in low infection intensity settings. An egg spiking study utilizing the same repeat-targeting assay described above determined that on average, DNA extraction from a sample containing a single *A. lumbricoides* egg gives a Cq value of approximately 24, with the most efficient extractions commonly producing Cq values in the range of 19-22 (32). This implies that efficiently extracted sub-egg quantities of target DNA should result in Cq values greater than this range. The bimodal distribution of *A. lumbricoides* Cq values likely reflects the presence of target sequences from samples with intact eggs and sub-egg level DNA (Fig 4). This distribution was not entirely unexpected as similar distributions have previously been described for other STH assays (18). The initial peak (near Cq=9) may represent signal derived from intact eggs, including samples with egg counts ranging from a single egg to many thousands of eggs. The second peak (near Cq=25) likely represents signal derived from sub-single egg quantities of DNA. Given that the target of this qPCR assay is estimated to number in the hundreds of thousands of copies per haploid genome, this distribution likely represents the detection of free DNA at various concentrations, with the upper tail representing concentrations approaching those found within a single intact egg and the lower tail representing concentrations of target approaching the limits of detection. Overall, 45% of positive *A. lumbricoides* samples had Cq values >22, suggesting that sub-egg quantities of DNA were common in our study population. The absence of discrete STH eggs in a stool sample does not necessarily imply that an individual is uninfected, and sub-egg level quantities of DNA could reflect clinical STH infection. There is large variability in the number of STH eggs secreted in stool, and eggs are not distributed evenly within stool (61,62). As intervention efforts continue to reduce infection rates and intensities, maximizing sensitivity of detection will play an increasingly important role in the programmatic decision-making process.

Prior studies have reported moderate-to-high correlations between epg and DNA concentration for all three STH (17–19,22,63), and one study reported that Cq values correlated well with expelled adult worms following anthelminthic treatment (22). We found moderate correlations between epg and Cq values that were lower than the correlations reported in prior studies (18,19,22). The likely misclassification of *A. lumbricoides* during Kato-Katz may have impacted estimates of epg in positive samples, resulting in overestimation in some samples and underestimation in others. This misclassification likely lowered the correlation between epg and Cq values. The strength of the correlation between epg and Cq values informs whether Cq value thresholds can be defined for STH infection intensity as is done using Kato-Katz (64). *A. lumbricoides* was the only STH in this study with a sufficiently large number of samples classified as moderate-to-heavy intensity using Kato-Katz. The majority of moderate-to-heavy intensity infections had Cq values of 10 or lower (Fig 4). However, because of the misclassification of *A. lumbricoides*, we interpret this pattern with caution.

Though a common critique of qPCR is its higher cost relative to Kato-Katz, comparisons of the materials required for each method suggest that costs for multi-parallel qPCR can be lower than those for Kato-Katz (19). As pointed out by Turner et al., even if qPCR is more costly than Kato-Katz, in an elimination setting, the continued use of low sensitivity diagnostics may impede efforts to determine when STH transmission has been interrupted which may unnecessarily prolong mass deworming administration. The cost of prolonged MDA is likely to exceed that of a more costly diagnostic (65).

## Conclusion

In a setting of STH elimination, highly sensitive diagnostics are needed to detect low intensity infections prior to elimination and to detect resurgent infections. Our results further support the recommendation that qPCR be considered the new gold standard STH diagnostic (22). Importantly, using qPCR instead of Kato-Katz in intervention trials can also reduce misclassification and bias of intervention effect estimates (53). Taken together, our findings underscore that molecular methods of detecting STH infection are more appropriate than copromicroscopic methods in settings with predominantly low infection intensity or with a goal to interrupt STH transmission.

## Supporting Information Legends

**S1 Appendix.** STARD Checklist

**S2 Appendix.** Bayesian latent class model specification

**S3 Appendix.** Bayesian latent class models priors and sensitivity analyses

**S4 Appendix.** Participant flow diagram

**S5 Appendix.** Taxonomic assignments for OTUs resulting from sequencing analysis with the Qiime2 pipeline. Following de-multiplexing by barcode, OTUs underwent taxonomic assignment. For each sample, the number of interlaced read pairs mapping to each taxonomic assignment is provided.

## Acknowledgements

This study was supported by an award from the Task Force for Global Health (NTD-SUSTAIN4 089G).

## Author contributions

**Conceptualization:** Jade Benjamin-Chung, Nils Pilotte, Ayse Ercumen, Benjamin F. Arnold, Steven A. Williams, Stephen P. Luby, John M. Colford, Jr.

**Data curation:** Jade Benjamin-Chung, Ayse Ercumen, Nils Pilotte

**Formal analysis:** Jade Benjamin-Chung, Nils Pilotte

**Funding acquisition:** Jade Benjamin-Chung, Benjamin F. Arnold, John M. Colford, Jr., Mahbubur Rahman, Stephen P. Luby, Steven A. Williams

**Investigation:** Jade Benjamin-Chung, Ayse Ercumen, Nils Pilotte, Jessica R. Grant, Jacqueline R.M.A. Maasch, Andrew M. Gonzalez, Brian P. Abrams and Ashanta C. Ester

**Methodology:** Jade Benjamin-Chung, Benjamin F. Arnold, Alan E Hubbard, Nils Pilotte, Steven A. Williams

**Project administration:** Jade Benjamin-Chung, Nils Pilotte

**Resources:** Ayse Ercumen, Mahbubur Rahman, Nils Pilotte, Steven A. Williams

**Software:** Jade Benjamin-Chung, Jessica R. Grant, Nils Pilotte

**Supervision:** Jade Benjamin-Chung, John M. Colford, Jr, Mahbubur Rahman, Nils Pilotte, Steven A. Williams

**Validation:** Ayse Ercumen, Rashidul Haque, Jacqueline R.M.A. Maasch, Andrew M. Gonzalez

**Visualization:** Jade Benjamin-Chung, Nils Pilotte

**Writing – original draft preparation:** Jade Benjamin-Chung, Nils Pilotte

**Writing – review and editing:** Jade Benjamin-Chung, Nils Pilotte, Jessica R. Grant, Jacqueline R.M.A. Maasch, Andrew M. Gonzalez, Brian P. Abrams, Ayse Ercumen, Benjamin F. Arnold, Mahbubur Rahman, Rashidul Haque, Alan E. Hubbard, Stephen P. Luby, Steven A. Williams, John M. Colford, Jr.

