## Supplementary material for "Comparison of multi-parallel qPCR and Kato-Katz for detection of soil-transmitted helminth infection among children in rural Bangladesh": S1 Appendix

**S1 Appendix: STARD Checklist**

| Section & Topic | No | Item | Reported on page # |
|---|---|---|---|
| **TITLE OR ABSTRACT** | | | |
| | **1** | Identification as a study of diagnostic accuracy using at least one measure of accuracy (such as sensitivity, specificity, predictive values, or AUC) | Abstract |
| **ABSTRACT** | | | |
| | **2** | Structured summary of study design, methods, results, and conclusions (for specific guidance, see STARD for Abstracts) | Abstract |
| **INTRODUCTION** | | | |
| | **3** | Scientific and clinical background, including the intended use and clinical role of the index test | Introduction, paragraph 2-4 |
| | **4** | Study objectives and hypotheses | Introduction, paragraph 5 |
| **METHODS** | | | |
| *Study design* | **5** | Whether data collection was planned before the index test and reference standard were performed (prospective study) or after (retrospective study) | Methods, paragraph 1-4 |
| *Participants* | **6** | Eligibility criteria | Methods, paragraph 1 |
| | **7** | On what basis potentially eligible participants were identified (such as symptoms, results from previous tests, inclusion in registry) | Methods, paragraph 1 |
| | **8** | Where and when potentially eligible participants were identified (setting, location and dates) | Methods, paragraph 1 |
| | **9** | Whether participants formed a consecutive, random or convenience series | Methods, paragraph 1 |
| *Test methods* | **10a** | Index test, in sufficient detail to allow replication | Methods, paragraph 3 |
| | **10b** | Reference standard, in sufficient detail to allow replication | Methods, paragraph 4-7 |
| | **11** | Rationale for choosing the reference standard (if alternatives exist) | Introduction, paragraph 2-4 |
| | **12a** | Definition of and rationale for test positivity cut-offs or result categories of the index test, distinguishing pre-specified from exploratory | Methods, paragraph 3 |
| | **12b** | Definition of and rationale for test positivity cut-offs or result categories of the reference standard, distinguishing pre-specified from exploratory | Methods, paragraph 6 |
| | **13a** | Whether clinical information and reference standard results were available to the performers/readers of the index test | Methods, paragraph 3-4 |
| | **13b** | Whether clinical information and index test results were available to the assessors of the reference standard | Methods, paragraph 7 |
| *Analysis* | **14** | Methods for estimating or comparing measures of diagnostic accuracy | Methods, paragraph 11-13 |
| | **15** | How indeterminate index test or reference standard results were handled | Methods, paragraph 4, 6 |
| | **16** | How missing data on the index test and reference standard were handled | N/A |
| | **17** | Any analyses of variability in diagnostic accuracy, distinguishing pre-specified from exploratory | Methods, paragraph 11-13 |
| | **18** | Intended sample size and how it was determined | Methods, paragraph 1 |
| **RESULTS** | | | |
| *Participants* | **19** | Flow of participants, using a diagram | S3 Appendix |
| | **20** | Baseline demographic and clinical characteristics of participants | Methods, paragraph 7; Results, paragraph 1 |
| | **21a** | Distribution of severity of disease in those with the target condition | Results, paragraph 2 |
| | **21b** | Distribution of alternative diagnoses in those without the target condition | N/A |
| | **22** | Time interval and any clinical interventions between index test and reference standard | Methods, paragraph 2 |

| | | | |
|---|---|---|---|
| *Test results* | **23** | Cross tabulation of the index test results (or their distribution) by the results of the reference standard | Results, paragraph 3 |
| | **24** | Estimates of diagnostic accuracy and their precision (such as 95% confidence intervals) | Results, paragraph 4 |
| | **25** | Any adverse events from performing the index test or the reference standard | N/A |
| **DISCUSSION** | | | |
| | **26** | Study limitations, including sources of potential bias, statistical uncertainty, and generalisability | Discussion, paragraph 3, 5 |
| | **27** | Implications for practice, including the intended use and clinical role of the index test | Discussion, paragraph 2, 3; Conclusion |
| **OTHER INFORMATION** | | | |
| | **28** | Registration number and name of registry | Methods, paragraph 1 |
| | **29** | Where the full study protocol can be accessed | Methods, paragraph 1 |
| | **30** | Sources of funding and other support; role of funders | Acknowledgements |
