## Supplementary material for "Comparison of multi-parallel qPCR and Kato-Katz for detection of soil-transmitted helminth infection among children in rural Bangladesh": S2 Appendix

### S2 Appendix: Bayesian latent class model specification

To estimate sensitivity and specificity of the STH diagnostic methods, we used Bayesian latent class analysis, which defines the true prevalence, sensitivity, and specificity as latent variables that are estimated simultaneously from the data and assumes no gold standard. We defined a model that accounted for covariance between diagnostic methods since both diagnostic methods are dependent upon the same underlying biological process.

We define the following 2x2 table, which compares Kato-Katz and qPCR test results:

| | qPCR + | qPCR − | |
|---|---|---|---|
| Kato-Katz + | $X_{++}$ | $X_{+-}$ | $T_1^+$ |
| Kato-Katz − | $X_{-+}$ | $X_{--}$ | $T_1^-$ |
| | $T_2^+$ | $T_2^-$ | |

The number of individuals who tested positive or negative by each test are denoted by $T_1^+$, $T_1^-$, $T_2^+$, and $T_2^-$, and the true number who tested positive and negative by each test are denoted by $D^+$ and $D^-$. $\pi$ is the true prevalence in the population. $Se_1$ is the true sensitivity of Kato-Katz in the population; $Se_2$ is the true sensitivity of qPCR in the population. $Sp_1$ is the true specificity of Kato-Katz in the population; $Sp_2$ is the true specificity of qPCR in the population.

We assume that the joint distribution of the two tests is multinomial $(X_{++}, X_{+-}, X_{-+}, X_{--}) \sim \text{Multi}(p_{++}, p_{+-}, p_{-+}, p_{--}, N)$ with probabilities defined as follows. These probabilities account for covariance between diagnostic methods since both diagnostic methods are dependent upon the same underlying biological process.

- $p_{++} = P(T_1^+, T_2^+) = [Se_1 Se_2 + cov(D_{12}^+)]\pi + [(1 - Sp_1)(1 - Sp_2) + cov(D_{12}^-)](1 - \pi)$
- $p_{+-} = P(T_1^+, T_2^-) = [Se_1(Se_2 - 1) - cov(D_{12}^+)]\pi + [(1 - Sp_1)Sp_2 - cov(D_{12}^-)](1 - \pi)$
- $p_{-+} = P(T_1^-, T_2^+) = [(Se_1 - 1)Se_2 - cov(D_{12}^+)]\pi + [Sp_1(1 - Sp_2) - cov(D_{12}^-)](1 - \pi)$
- $p_{--} = P(T_1^-, T_2^-) = [(Se_1 - 1)(1 - Se_2) + cov(D_{12}^+)]\pi + [Sp_1 Sp_2 - cov(D_{12}^-)](1 - \pi)$

The conditional correlations between two test outcomes are calculated as follows:

Conditional correlations between two test outcomes for infected individuals: $\rho_{D+} = \frac{covD_{12}^+}{\sqrt{Se_i(1-Se_i)Se_j(1-Se_j)}}$

Conditional correlations between two test outcomes for non-infected individuals: $\rho_{D-} = \frac{covD_{12}^-}{\sqrt{Sp_i(1-Sp_i)Sp_j(1-Sp_j)}}$

Further, we assume the following bounds on the covariance:

- $(Sp_1 - 1)(1 - Sp_2) \leq cov(D_{12}^+) \leq \min(Sp_1, Sp_2) - Sp_1 Sp_2$
- $(Se_1 - 1)(1 - Se_2) \leq cov(D_{12}^-) \leq \min(Se_1, Se_2) - Se_1 Se_2$
