## Supplementary material for "Comparison of multi-parallel qPCR and Kato-Katz for detection of soil-transmitted helminth infection among children in rural Bangladesh": S3 Appendix

### S3 Appendix: Bayesian latent class models priors and sensitivity analyses

**Table 1. Prior distributions used in Bayesian latent class analysis models**

| | ***A. lumbricoides*** | **Hookworm** | ***T. trichiura*** |
|---|---|---|---|
| Prevalence | Beta distribution with shape parameters $\alpha$=1, $\beta$=1 | Beta distribution with shape parameters $\alpha$=1, $\beta$=1 | Beta distribution with shape parameters $\alpha$=1, $\beta$=1 |
| Kato-Katz sensitivity | Beta distribution with shape parameters $\alpha$=1, $\beta$=1 | Beta distribution with shape parameters $\alpha$=3, $\beta$=3 | Beta distribution with shape parameters $\alpha$=3, $\beta$=3 |
| Kato-Katz specificity | Beta distribution with shape parameters $\alpha$=1, $\beta$=1 | Uniform distribution with minimum=0.95, maximum=1 | Uniform distribution with minimum=0.95, maximum=1 |
| qPCR sensitivity | Uniform distribution with minimum=0.60, maximum=1 | Uniform distribution with minimum=0.80, maximum=1 | Uniform distribution with minimum=0.80, maximum=1 |
| qPCR specificity | Uniform distribution with minimum=0.95, maximum=1 | Uniform distribution with minimum=0.95, maximum=1 | Uniform distribution with minimum=0.95, maximum=1 |

Due to the discrepancy between Kato-Katz and qPCR results for *A. lumbricoides*, we chose non-informative prior distributions for Kato-Katz sensitivity and specificity and a less informative prior distribution for qPCR sensitivity.

**Table 2. Alternative prior distributions used in sensitivity analysis for Bayesian latent class analysis models for *A. lumbricoides***

| | **Sensitivity analysis 1**:<br><br>More informative prior for Kato-Katz sensitivity and specificity | **Sensitivity analysis 2:**<br><br>More informative prior for qPCR sensitivity | **Sensitivity analysis 3:**<br><br>More informative prior for sensitivity and specificity of both Kato-Katz and qPCR |
|---|---|---|---|
| Prevalence | Beta distribution with shape parameters $\alpha$=1, $\beta$=1 | Beta distribution with shape parameters $\alpha$=1, $\beta$=1 | Beta distribution with shape parameters $\alpha$=1, $\beta$=1 |
| Kato-Katz sensitivity | Beta distribution with shape parameters $\alpha$=3, $\beta$=3 | Beta distribution with shape parameters $\alpha$=3, $\beta$=3 | Beta distribution with shape parameters $\alpha$=3, $\beta$=3 |
| Kato-Katz specificity | Uniform distribution with minimum=.5, maximum=1 | Beta distribution with shape parameters $\alpha$=1, $\beta$=1 | Uniform distribution with minimum=.5, maximum=1 |
| qPCR sensitivity | Uniform distribution with minimum=0.6, maximum=1 | Uniform distribution with minimum=0.8, maximum=1 | Uniform distribution with minimum=0.8, maximum=1 |
| qPCR specificity | Uniform distribution with minimum=0.95, maximum=1 | Uniform distribution with minimum=0.95, maximum=1 | Uniform distribution with minimum=0.95, maximum=1 |

**Table 3. Sensitivity analysis using alternative prior distributions in Bayesian latent class analysis models for *A. lumbricoides***

| | | Kato-Katz | | qPCR | |
|---|---|---|---|---|---|
| **Analysis** | **Description of priors** | **Sensitivity (95% BCI)** | **Specificity (95% BCI)** | **Sensitivity (95% BCI)** | **Specificity (95% BCI)** |
| Primary analysis result | Noninformative priors | 49 (34, 64) | 68 (61, 77) | 79 (61, 99) | 97 (95, 100) |
| Sensitivity analysis 1 | More informative prior for Kato-Katz sensitivity and specificity | 49 (34, 64) | 68 (62, 76) | 80 (61, 99) | 97 (95, 100) |
| Sensitivity analysis 2 | More informative prior for qPCR sensitivity | 49 (38, 59) | 67 (63, 71) | 90 (80, 99) | 97 (95, 100) |
| Sensitivity analysis 3 | More informative prior for sensitivity and specificity of both Kato-Katz and qPCR | 49 (39, 59) | 67 (63, 71) | 90 (80, 99) | 97 (95, 100) |
