## Supplementary material for "Comparison of multi-parallel qPCR and Kato-Katz for detection of soil-transmitted helminth infection among children in rural Bangladesh": S4 Appendix

**S4 Appendix: Participant flow diagram**

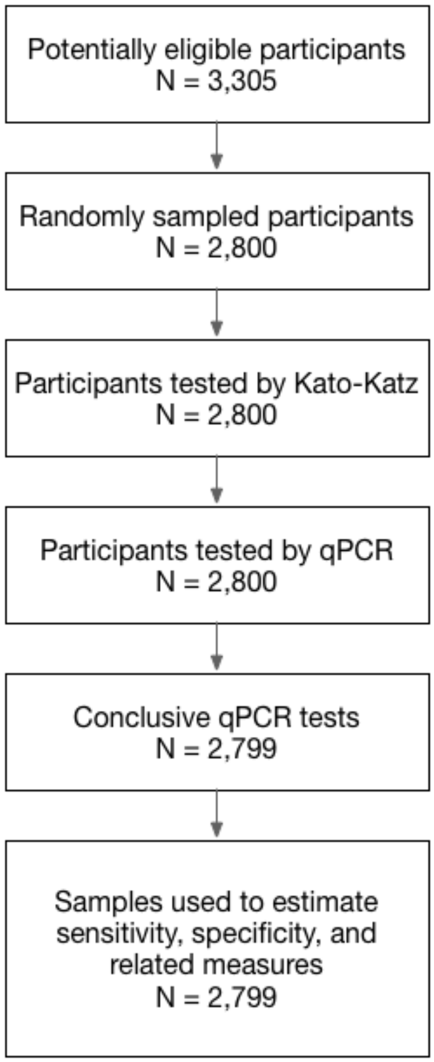
